# Collateral Sensitivity Associated with Antibiotic Resistance Plasmids

**DOI:** 10.1101/2020.07.10.198259

**Authors:** Cristina Herencias, Jerónimo Rodríguez-Beltrán, Ricardo León-Sampedro, Aida Alonso-del Valle, Jana Palkovičová, Rafael Cantón, Álvaro San Millán

## Abstract

Collateral sensitivity (CS) is a promising alternative approach to counteract the rising problem of antibiotic resistance (ABR). CS occurs when the acquisition of resistance to one antibiotic produces increased susceptibility to a second antibiotic. For CS to be widely applicable in clinical practice, it would need to be effective against the different resistance mechanisms available to bacteria. Recent studies have focused on CS strategies designed against ABR mediated by chromosomal mutations. However, one of the main drivers of ABR in clinically relevant bacteria is the horizontal transfer of ABR genes mediated by plasmids. Here, we report the first analysis of CS associated with the acquisition of complete ABR plasmids, including the clinically important carbapenem-resistance conjugative plasmid pOXA-48. In addition, we describe the conservation of CS in clinical *E. coli* isolates and its application to the selective elimination of plasmid-carrying bacteria. Our results provide new insights that establish the basis for developing CS-informed treatment strategies to combat plasmid-mediated ABR.

## Introduction

The rapid evolution of antibiotic resistance (ABR) in bacteria reduces the utility of clinically relevant antibiotics, making ABR one of the major challenges facing public health (Jim, 2016; MacLean and San Millan, 2019). There is therefore an urgent need for new treatment strategies to fight against resistant pathogens (Baym et al., 2016). Among the most promising alternatives, much attention has focused on collateral sensitivity (CS). CS occurs when the acquisition of resistance to one antibiotic causes increased susceptibility to a second antibiotic (Szybalski and Bryson, 1952) and can be exploited for the design of multi-drug strategies that select against ABR (Imamovic and Sommer, 2013; Lázár et al., 2013). Moreover, recent evidence suggests that the development of resistance might be prevented by treatments based on the sequential cycling of antibiotics whose resistance mechanisms produce reciprocal CS (Imamovic and Sommer, 2013). Several studies have cataloged CS networks emerging from ABR mutations in chromosomal genes (Imamovic and Sommer, 2013; Lázár et al., 2013; Maltas and Wood, 2019; Podnecky et al., 2018; Roemhild et al., 2020), but the CS effects produced by the horizontal acquisition of ABR plasmids remain poorly understood.

Mobile genetic elements, especially plasmids, play a crucial role in the dissemination of ABR genes between clinical pathogens and are one of the major drivers of the alarming worldwide rise in ABR (Partridge et al., 2018). Despite the evolutionary advantage conferred by plasmids in the presence of antibiotics, plasmids acquisition tends to produce common metabolic alterations in the host bacterium (San Millan et al., 2018), that typically translate into a fitness cost (San Millan and MacLean, 2017). We reasoned that the physiological alterations produced upon acquisition of ABR plasmids might lead to exploitable CS responses. To test this hypothesis, we analyzed the collateral effects associated with the acquisition of 6 natural plasmids carrying clinically relevant ABR genes in *Escherichia coli* MG1655. Our results reveal that most ABR plasmids, including the clinically important carbapenem-resistance conjugative plasmid pOXA-48, produce CS events of moderate effect. To extend our results, we explored the degree of conservation of CS responses associated with pOXA-48 across phylogenetically diverse wild-type *E. coli*. Our results demonstrate that these responses can be exploited to preferentially eradicate plasmid-carrying bacteria.

## Results and discussion

### Collateral sensitivity induced by ABR plasmids

To test whether ABR plasmids elicit exploitable CS to other antibiotics, we selected 6 clinically relevant ABR plasmids (see methods, Table 1 and Supplementary Figure 1). All plasmids carried important β-lactamase genes, including extended-spectrum β-lactamases (ESBLs), AmpC-type β-lactamases, or carbapenemases, as well as other relevant ABR genes (Table 1). These plasmids belonged to a broad range of incompatibility groups, covered a range of different sizes, included conjugative and non-conjugative specimens, and produced variable fitness effects, ranging from 10% to 27% reductions in relative fitness, when introduced into *E. coli* MG1655 (Figure 1A).

**Figure 1.**
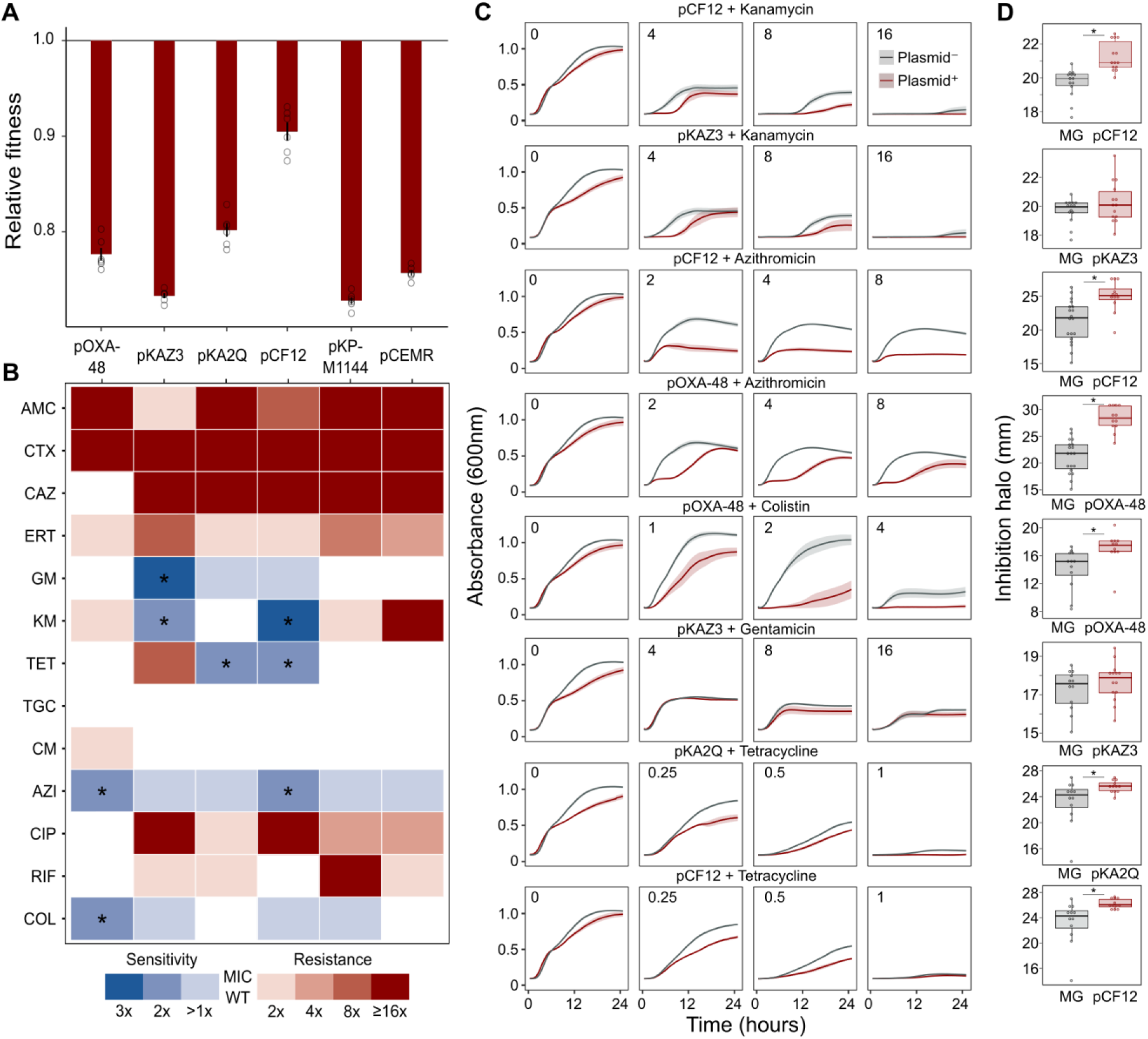
Collateral sensitivity and fitness effects associated with ABR plasmids. (**A**) Competition-assay–determined fitness of plasmid-carrying clones relative to the plasmid-free MG1655 strain. Bars represent the means of 6 independent experiments (represented by dots), and error bars represent the standard error of the mean. (**B**) Heat-map representing collateral responses to antibiotics associated with plasmid acquisition. The color code represents the fold change of MIC in plasmid-carrying derivatives compared to plasmid-free MG1655 (see legend); MIC values were the median of 4-5 independent determinations. Asterisks indicate a ≥ 2-fold decrease in MIC, which we defined as significant CS. AMC, amoxicillin-clavulanic acid; CTX, cefotaxime; CAZ, ceftazidime; ERT, ertapenem; GM, gentamicin; KM, kanamycin; TET, tetracycline; TGC, tigecycline; CM, chloramphenicol; AZI, azithromycin; CIP, ciprofloxacin; RIF, rifampicin; and COL, colistin. (**C**) Growth curves of MG1655 and plasmid-carrying MG1655 strains exposed to increasing antibiotic concentrations. The antibiotic concentration, in mg/L, is indicated at the top left corner of each panel. Lines represent the mean of 6 biological replicates, and the shaded area indicates standard error of the mean. (**D**) Boxplot representations of the inhibition halo diameters, in mm, obtained from disk-diffusion antibiograms of plasmid-free and plasmid-carrying MG1655. Horizontal lines within boxes indicate median values, upper and lower hinges correspond to the 25th and 75th percentiles, and whiskers extend to observations within 1.5 times the interquartile range. Individual data points are also represented (11-15 biological replicates). Asterisks indicate statistically significant differences (unpaired t-test with Welch’s correction *P*< 0.044).

**Table 1.**
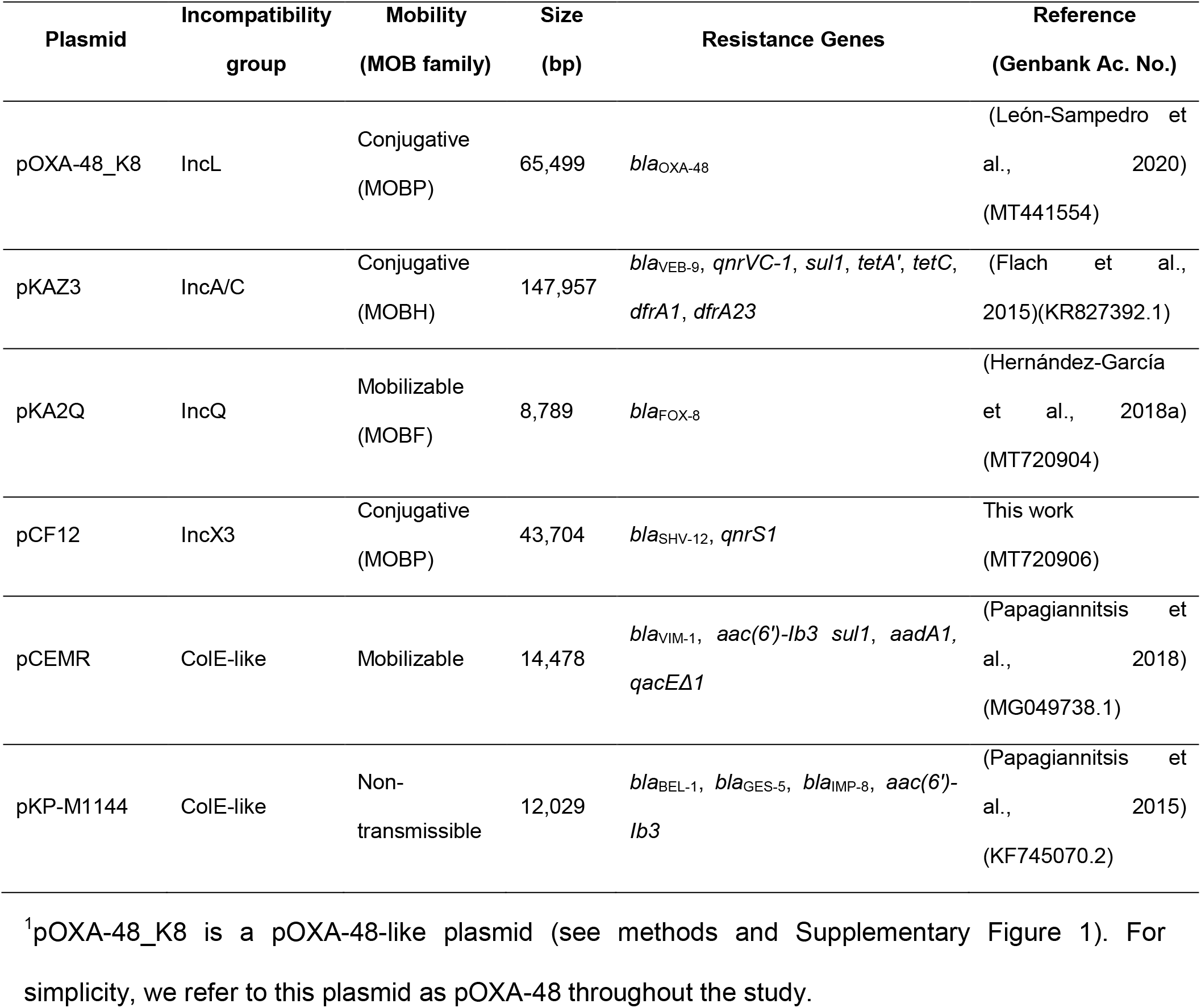
Plasmids used in this study.

We performed dose-response experiments with 13 antibiotics from 8 drug families (Supplementary Table 1). For this analysis, we determined the minimal inhibitory concentration (MIC, see methods) using the broth microdilution method for plasmid-free MG1655 and each of its 6 plasmid-carrying derivatives (Supplementary Table 1). We analyzed the collateral antibiotic susceptibility effects associated with the presence of each plasmid, measured as the fold-change in the antibiotic MIC between plasmid-carrying and plasmid-free bacteria (Figure 1B). As expected from their genetic content, all plasmids conferred resistance to β-lactam antibiotics and most of them conferred additional resistance to other antibiotics from unrelated families (Figure 1B and Supplementary Table 1). CS was defined as a minimum 2-fold reduction in MIC (calculated as the fold-change of the median MIC value obtained from 4-5 independent MIC determinations). Using this definition, we identified 8 instances of plasmid-induced CS. The plasmids that produced CS effects were pOXA-48 (to colistin and azithromycin), pKAZ3 (to kanamycin and gentamicin), pKA2Q (to tetracycline), and pCF12 (to kanamycin, azithromycin, and tetracycline).

To validate the MIC results, we conducted two further series of experiments for the 8 cases of plasmid-induced CS: (i) complete growth curves under a range of antibiotic concentrations and (ii) disk-diffusion antibiograms on agar plates. Analysis of the area under the growth curve, which integrates all growth parameters (maximum growth rate, lag duration and carrying capacity) (DelaFuente et al., 2020), confirmed significant plasmid-mediated CS effects at different antibiotic concentrations for 6 of the combinations (pOXA-48–azithromycin, pCF12–azithromycin, pOXA-48–colistin, pCF12–kanamycin, pKAZ3–kanamycin, and pKA2Q–tetracycline; Figure 1C, Supplementary Figure 2 and Supplementary Table 2). The disk-diffusion results largely agreed with the MIC and growth-curve data, showing a general trend towards an increased diameter of the antibiotic inhibition zone in plasmid-carrying strains compared with plasmid-free MG1655 (Figure 1D). These differences were statistically significant for pCF12-kanamycin, pCF12-azithromycin, pOXA-48-azithromycin, pOXA-48-colistin, pCF12-tetracycline, and pKA2Q-tetracycline (unpaired t-test with Welch’s correction; P<0.044 in all cases). Overall, 7 of the 8 CS instances identified by MIC testing were validated with at least one of the additional methods (growth curves or disk-diffusion assay), and 6 of them showed full agreement across all 3 methods. The disk-diffusion assays produced notably robust results with minimum hands-on time, suggesting that this technique is appropriate for screening CS responses in large strain collections. Together, these results demonstrate that ABR plasmids produce moderate but significant CS to different antibiotics, comparable to that generated by ABR chromosomal mutations in *E. coli* (Imamovic and Sommer, 2013; Lázár et al., 2013; Podnecky et al., 2018).

### Conservation of pOXA-48–mediated CS

The success and applicability of CS-informed therapeutic strategies are crucially dependent on the conservation of CS across diverse genetic contexts (Podnecky et al., 2018). To address the phylogenetic preservation of plasmid-induced CS, we focused on the plasmid pOXA-48, which confers resistance to last-resort carbapenem antibiotics, and whose prevalence in clinical settings is rising alarmingly (David et al., 2019). We tested the degree of conservation of pOXA-48–mediated CS patterns (to azithromycin and colistin) across phylogenetically diverse *E. coli* strains using disk-diffusion assays. We determined antibiotic susceptibility in 9 diverse *E. coli* clinical isolates (Figure 2A) and their pOXA-48–carrying derivatives. CS to azithromycin showed a striking degree of conservation across the strains, with 8 out of 9 plasmid-carriers showing significantly higher susceptibility than their plasmid-free counterparts. In contrast, CS to colistin was conserved in only 2 of the tested strains (Figure 2B). Nevertheless, all strains showed CS responses to at least one antibiotic, suggesting that pOXA-48 elicits CS responses that could potentially be exploited therapeutically.

**Figure 2.**
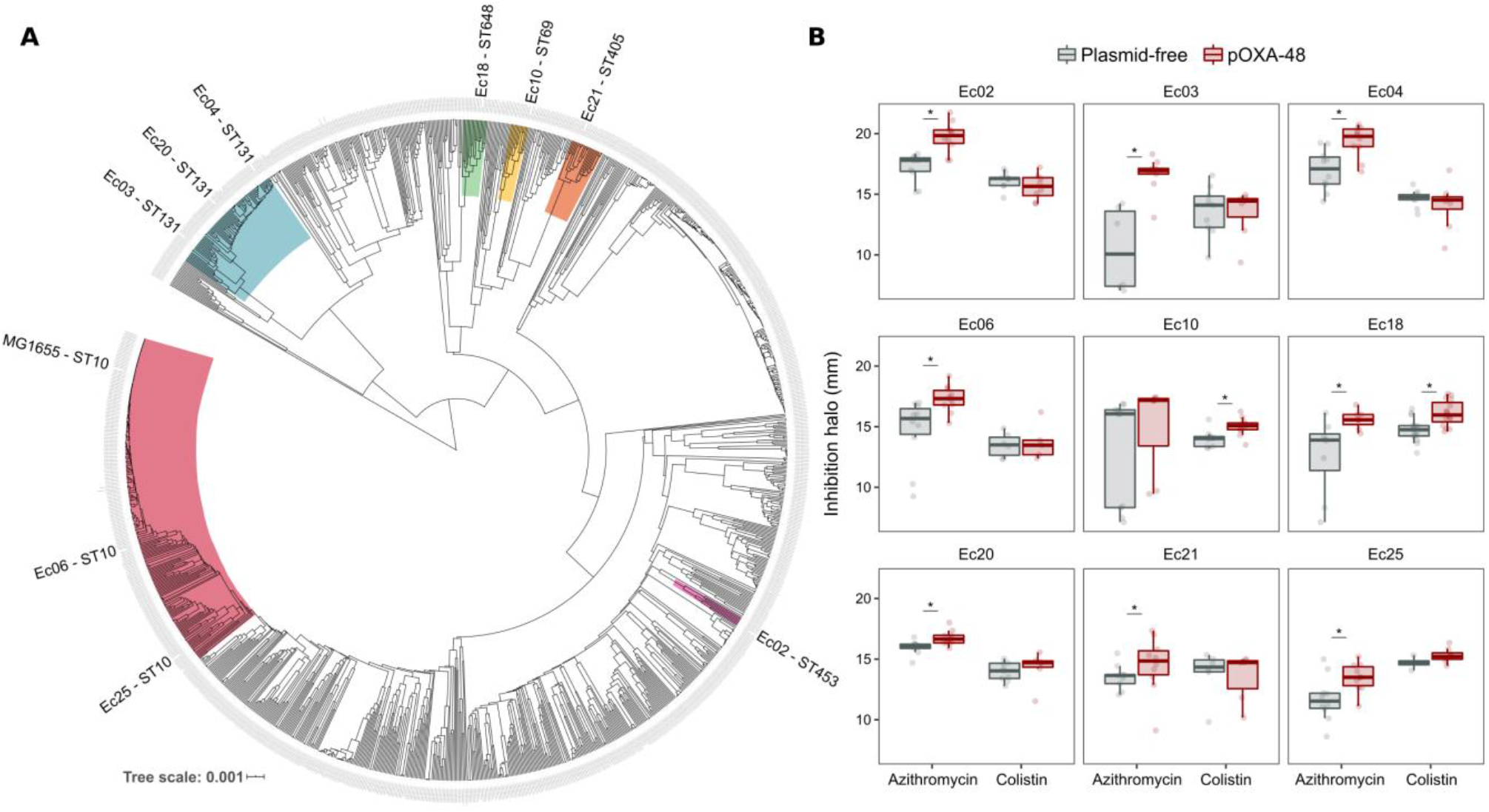
Phylogeny and collateral sensitivity profiles of *E. coli* clinical isolates. (**A**) Phylogenetic tree of *E. coli* species, highlighting the strains used in this study and their sequence types (ST). Colors within tree branches represent different STs. The tree depicts the phylogenetic relationships of 1344 representative *E. coli* genomes obtained from NCBI. (**B**) Representation of the inhibition halo diameters, in mm, obtained from disk-diffusion antibiograms of clinical isolates and their transconjugants carrying pOXA-48. Horizontal lines inside boxes indicate median values, upper and lower hinges correspond to the 25th and 75th percentiles, and whiskers extend to observations within 1.5 times the interquartile range. Individual data points are also represented (6-12 replicates, jittered to facilitate data visualization). Asterisks denote statistically significant differences (t-test with Welch’s correction P<0.035 in all cases).

### Exploitation of CS for the preferential extinction of pOXA-48–carrying *E. coli*

We next tested the potential of exploiting CS effects to eradicate plasmid-carrying subpopulations. We rationally selected 4 *E. coli* isolates and their pOXA-48 carrying derivatives with defined CS profiles: to colistin only (Ec10), to azithromycin only (Ec21 and Ec25), or to both antibiotics (Ec18). MG1655 was included as a control. Each bacterial population was propagated in the presence of a single antibiotic or with sequential exposure to both antibiotics over 2 days. On the first day of the experiment, 24 bacterial populations of each strain were inoculated into media containing either colistin (4 mg/L) or azithromycin (8 mg/L). After growth for 22 hours, surviving populations were transferred to fresh media containing colistin or azithromycin in a full factorial experimental design. This approach gave rise to 4 antibiotic treatments: 2 single-drug treatments (denoted Azi→ Azi and Col→ Col) and 2 treatments in which the antibiotic changed (Azi→ Col and Col→ Azi). In 11 of the 20 possible strain-treatment combinations, survival patterns showed significantly higher mortality rates for the pOXA-48–carrying strains than for their plasmid-free counterparts (Figure 3; log-rank test P<0.001 in all cases). Crucially, the mortality patterns observed were consistent with the CS patterns obtained in the disk-diffusion technique (Figure 2B). For instance, plasmid-carrying MG1655 and Ec18 strains, which show CS to both antibiotics, exhibited significant reductions in survival in all treatments. In contrast, for strain Ec10, which shows CS exclusively to colistin, significant extinction was observed only when the treatment included colistin as the first anti-biotic (Col→ Col or Col→ Azi).

**Figure 3.**
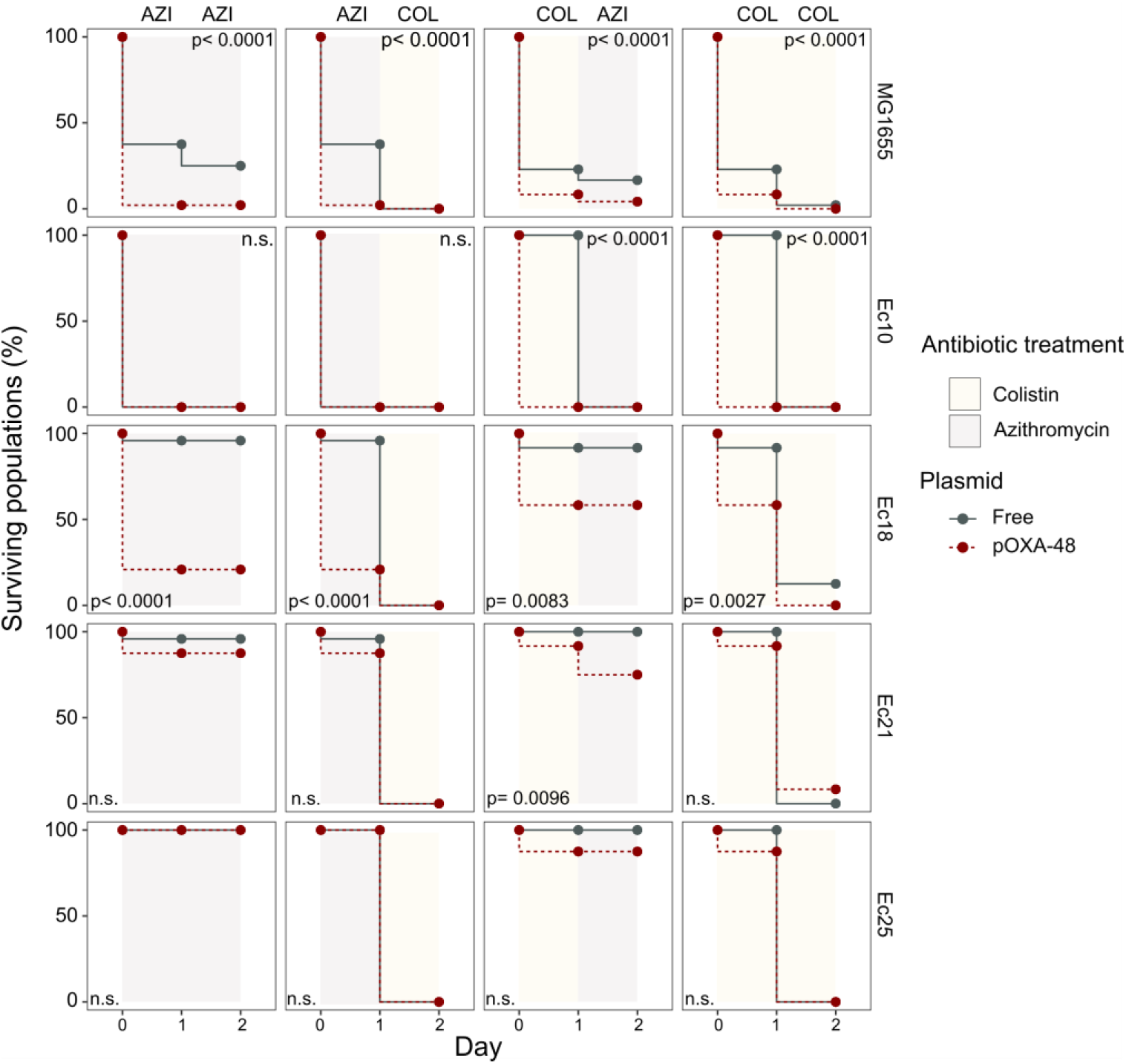
Plasmid-mediated CS can be exploited to preferentially eliminate plasmid-carrying bacteria. Survival curves of plasmid-free (gray solid lines) and plasmid-carrying (red dashed lines) bacterial populations in response to constant or sequential antibiotic treatments. Bacteria were cultured for 2 days with either constant antibiotic exposure or with a change of antibiotic after the first day; treatments are specified by the background colors in each panel. Log-rank test p-values assessing statistically different survival rates are shown. n.s. non-significant. For strains Ec10, Ec18, Ec21 and Ec25 n=24; for MG1665 n=48. AZI, azithromycin; COL, colistin.

### Concluding remarks

Our results reveal that the acquisition of clinically-relevant ABR plasmids induces CS to unrelated antibiotics. These findings thus serve as a stepping stone toward the development of new approaches aimed at blocking plasmid-mediated horizontal spread of ABR genes. These antiplasmid strategies could help to tackle the alarming clinical and community spread of ABR. However, the molecular basis of plasmid-associated CS is currently unknown. The feasibility of rational broad-spectrum anti-plasmid strategies will depend on gaining comprehensive knowledge about the molecular mechanisms that increase antibiotic susceptibility upon plasmid acquisition. Until these therapies are available, CS-informed empirical treatment of plasmid-carrying bacteria has the potential to help resolve antibiotic resistant infections. In this regard, our results suggest that colistin and azithromycin may be valuable antibiotics for the treatment of pOXA-48–carrying enterobacteria. Both antibiotics are already used to treat enterobacterial infections (Lübbert, 2016; Morrill et al., 2015), and colistin in particular is currently used as a last resort antibiotic against carbapenem-resistant enterobacteria (Rodriguez-Baño et al., 2018). In summary, the findings presented here open new avenues of research aimed at understanding plasmid-associated CS and pave the way for the development of new CS-based treatment strategies to combat plasmid-mediated ABR.

## Material and Methods

### Bacterial strains, plasmids and culture conditions

All experiments were performed in liquid Lennox lysogeny broth (LB; CONDA) or on LB agar (15 g/L) unless indicated. Mueller Hinton II broth (Oxoid) was used to verify that MIC values of *E. coli* MG1655 were comparable with those obtained in LB medium. The antibiotics used were amoxicillin-clavulanic acid, ceftazidime, cefotaxime, gentamicin (Normon), ertapenem (Merck Sharp & Dohme), kanamycin, chloramphenicol, tetracycline (Nzytech), tigecycline (Pfizer), azithromycin, colistin (Altan Pharmaceuticals), and rifampicin (Sanofi-aventis).

Plasmids pKAZ3 (Accession Number KR827392.1), pKP-M1144 (KF745070.2), pKA2Q (MT720904) and pEncl-30969cz (MG049738.1) were previously published (Flach et al., 2015; Hernández-García et al., 2018a; Papagiannitsis et al., 2015, 2018). Plasmid pCEMR is a derivative of pEncl-30969cz in which a spontaneous deletion of ~19kb had occurred (MT720903). Plasmids pOXA-48_K8 (MT441554) and pCF12 (MT720906) were isolated from rectal swabs of patients hospitalized in the Ramón y Cajal University Hospital in the framework of the R-GNOSIS project (Hernández-García et al., 2018b). pOXA-48_K8 is a pOXA-48-like plasmid that shares a 100% coverage and >99% identity with the first described pOXA-48 plasmid (León-Sampedro et al., 2020; Poirel et al., 2012). For simplicity, we refer to this plasmid as pOXA-48 throughout the text. Plasmids pKAZ3, pOXA-48 and pCF12 were conjugated into MG1655 using *E. coli* β3914 as donor strain, which is auxotrophic for diaminopimelic acid (Le Roux et al., 2007). Transconjugants were selected in LB plates containing ampicillin (100 mg/L) for plasmids pKAZ3 and pOXA-48 or aztreonam (25 mg/L) for pCF12 plasmid. Plasmids pCEMR, pKP-M1144 and pKA2Q were purified using a commercial mini-prep kit (Macherey-Nagel) and transformed into calcium chloride competent MG1655 cells. Transformant cells were selected on LB-ampicillin plates (100 mg/L). Strain constructions were verified by assessing that the antibiotic resistance profile of plasmid-bearing MG1655 matched the expected resistance of the plasmid (Supplementary Table 1) and by sequencing the complete genomes of transconjugant clones, which also revealed that no significant chromosomal mutations accumulated during the process of plasmid acquisition (0-2 mutations between plasmid-carrying and plasmid-free complete genomes, see Supplementary Table 3).

The clinical *E. coli* strains used in this study were isolated during the R-GNOSIS project, which included representative ESBL-producing enterobacteria from hospitalized patients in the Ramon y Cajal University Hospital during a 2 year period (Hernández-García et al., 2018b; León-Sampedro et al., 2020). The *E. coli* strains used here, and their corresponding transconjugants, were described and characterized previously (Alonso-del Valle et al., 2020) and Supplementary Table 4).

### Antimicrobial susceptibility testing

Single colonies of plasmid-free and plasmid-carrying bacteria were inoculated in LB starter cultures and incubated at 37 °C for 16 hours at 225 rpm (4-5 biological replicates performed on different days). Each culture was diluted 1:2,000 in LB medium (~5·10^4^ colony forming units) and 200 μl were added to a 96-well microtiter plate containing the appropriate antibiotic concentration. MIC values were measured after 22 hours of incubation at 37 °C. Optical density at 600 nm (OD_600_) was determined in a Synergy HTX (BioTek) plate reader after 30 seconds of orbital shaking. MIC values corresponded with the lowest antibiotic concentration that resulted in OD_600_< 0.2. We found that this threshold is qualitatively consistent with standard practice (i.e. visual determination of growth inhibition), while providing higher accuracy and repeatability. The final MIC value is the median of 4 to 5 replicates. MIC determinations of plasmid-free and plasmid-carrying strains were performed in parallel to ensure reproducibility. Fold change in MIC was determined as the ratio between the median MIC form plasmid-carrying derivatives and that form the wild-type strain.

### Disk-diffusion susceptibility testing

LB plates were swabbed with a 0.5 McFarland matched bacterial suspension and sterile disks containing the antibiotic were placed on the plates. Plates were incubated at 37 °C for 22 hours and pictures were taken using an in-house photographic system and a cell phone (Huawei Mate 20). Inhibition halos were measured using ImageJ software. Antibiotic disk content used in diffusion assays were as follows (all from Bio-Rad): gentamicin 10 μg, azithromycin 15 μg, kanamycin 30 μg, colistin 10 μg, and tetracycline 30 μg.

### Bacterial growth curves

Starter cultures were prepared and incubated as described above. Each culture was diluted 1:2,000 in LB medium and 200 μl were added to a 96-well microtiter plate containing the appropriate antibiotic concentration. Plates were incubated 24 hours at 37 °C with strong orbital shaking before reading OD_600_ every 10 minutes in a Synergy HTX (BioTek) plate reader. 6 biological replicates were performed for each growth curve. The area under the growth curve was obtained using the *‘auc’* function from the *‘flux’* R package. Data was represented using a R custom script and the *‘ggplot2’* package.

### Competition assays

Competition assays were performed to measure the relative fitness of plasmid-carrying strains using flow cytometry as previously described (Rodriguez-Beltran et al., 2018). Briefly, competitions were performed between MG1655 clones carrying each of the 6 plasmids and plasmid-free MG1655 against a MG1655 derivative carrying an arabinose inducible chromosomal copy of *gfp* (MG1655::*gfp*). Pre-cultures were incubated at 37 °C with 225 rpm shaking overnight in 96-well plates carrying 200 μl of LB broth per well. Pre-cultures were mixed at 1:1 proportion and diluted 1:400 in fresh media. Initial proportions of GFP and non-fluorescent competitors were confirmed in a CytoFLEX Platform (Beckman Coulter Life Sciences) flow cytometer, recording 10,000 events per sample. To measure these proportions, we incubated a culture aliquot in NaCl 0.9% containing 0.5% L-arabinose for 1.5 hours to induce the expression of the chromosomal GFP. Mixtures were competed for 22 hours in LB medium at 37 °C with shaking (225 rpm). Final proportions were estimated again by flow cytometry as described above. The fitness of each strain relative to MG1655::*gfp* was calculated using the formula: *w*=ln(N_final,gfp^-^_/N_*initial,gfp^-^*_)/ln(N_*final,gfp^+^*_/N_*initial,gfp^+^*_) where *w* is the relative fitness of the non GFP-tagged strain, N_*initial,gfp^-^*_ and N_*final,gfp^-^*_ are the numbers of non GFP-tagged cells before and after the competition and *N_initial,gfp_*^+^ and *N_final,gfp+_* are the numbers of MG1655::*gfp* cells before and after the competition. Data was normalized by dividing the relative fitness of plasmid-carrying derivative strains by the fitness calculated for plasmid-free MG1655. Six biological replicates were performed for each competition.

### CS-informed antibiotic treatments

Overnight bacterial cultures were diluted and seeded into 24 independent wells of a 96-well plate filled with 200 μl of LB. After 16 hours of incubation at 37 °C, these populations were diluted 1:2,000 into fresh medium containing either azithromycin (8 mg/L) or colistin (4 mg/L) and grown at 37 °C for 22 hours. The following day, bacterial populations were diluted (1:2,000) and inoculated into new plates containing, again, either azithromycin or colistin and allowed to grow as above. This approach led to four antibiotic treatments, two in which the antibiotic remains constant (Azi→ Azi and Col→ Col) and two in which the antibiotic treatment alternates (Azi→ Col and Col→ Azi). The survival of bacterial populations was assessed by measuring OD_600_ every day. Populations that did not reach an OD_600_ of 0.2 were declared extinct.

### Whole genome sequencing

Genomic DNA was isolated using the Wizard genomic DNA purification kit (Promega) and quantified using the QuantiFluor dsDNA system (Promega) following manufacturer’s instructions. Whole genome sequencing was conducted at the Wellcome Trust Centre for Human Genetics (Oxford, UK), using the Illumina HiSeq4000 platform with 125 base pair (bp) paired-end reads. For plasmid pCF12, first described in this study, we performed additional long-read sequencing using PacBio technologies. PacBio sequencing was performed at The Norwegian Sequencing Centre PacBio RSII platform using P5-C3 chemistry. Illumina and PacBio sequence reads were trimmed using the Trimmomatic v0.33 tool (Bolger et al., 2014). SPAdes v3.9.0 was used to generate *de novo* assemblies from the trimmed sequence reads with the −cov-cutoff flag set to ‘auto’ (Bankevich et al., 2012). Unicycler was used to generate hybrid assemblies from Illumina and PacBio data (Wick et al., 2017). QUAST v4.6.0 was used to generate assembly statistics (Gurevich et al., 2013). All the *de novo* assemblies reached enough quality including total size of ~4.6 Mb, and a total number of contigs over 1 kb lower than 200. Prokka was used to annotate the *de novo* assemblies (Seemann, 2014). The plasmid content of each genome was characterised using PlasmidFinder 2.1 (Carattoli et al., 2014), and the antibiotic resistance gene content was characterised with ResFinder 3.2 (Zankari et al., 2012). MOB families were characterised using MOB-typer tool included in MOB-suite (Robertson and Nash, 2018). Variant calling was performed using Snippy v4.6.0 (https://github.com/tseemann/snippy). The sequences generated and analysed during the current study are available in the Sequence Read Archive (SRA) repository, BioProject ID: PRJNA644278 (https://www.ncbi.nlm.nih.gov/bioproject/644278), and Genbank accession numbers are provided in Table 1.

To determine the distribution of the *E. coli* isolates across the phylogeny of the species, we obtained 1334 assemblies of *Escherichia coli* complete genomes from the RefSeq database (https://www.ncbi.nlm.nih.gov/assembly). Distances between genomes were established using Mash v2.0 (Ondov et al., 2016) and a phylogeny was constructed with mashtree v0.33 (Katz et al., 2019). The tree was represented with midpoint root using the phylotools package in R (https://github.com/helixcn/) and visualised using the iTOL tool (Letunic and Bork, 2011). To determine the phylotype of each genome, we used the ClermonTyping tool (Beghain et al., 2018).

### Statistical analyses

Data sets were analyzed using Prism 6 software (GraphPad Software Inc.) and R. Normality of the data was assessed by visual inspection and the Shapiro-Wilk test. ANOVA test were perfomed to ascertain the effect of the ‘plasmid x antibiotic concentration’ interaction term in the analysis of growth curves (area under the growth curve). Statistical analyses of the disk-diffusion halos were performed using unpaired t-tests with Welch’s correction. Survival curves were analyzed using the log-rank test within the ‘*survminer*’ R package.

## Acknowledgments

We are grateful to thank Costas Papagiannitsis, Carl-Frederik Flach, Fabrice Elia Graf and Martin Palm for generous gifts of plasmids. We also appreciate the technical support of Carmen de la Vega and Laura Jaraba. This work was supported by the European Research Council under the European Union’s Horizon 2020 research and innovation programme (ERC grant agreement no. 757440-PLASREVOLUTION) and by the *Instituto de Salud Carlos III* (grant PI16-00860) co-funded by European Development Regional Fund “a way to achieve Europe”. ASM is supported by a Miguel Servet Fellowship (MS15-00012). JRB is a recipient of a *Juan de la Cierva-Incorporación* Fellowship (IJC2018-035146-I) co-funded by *Agencia Estatal de Investigación del Ministerio de Ciencia e Innovación*. CH is supported by *Comunidad Autónoma de Madrid* (PEJD-2018-POST/BMD-8016).

## Supplementary Material

**Supplementary Figure 1.**
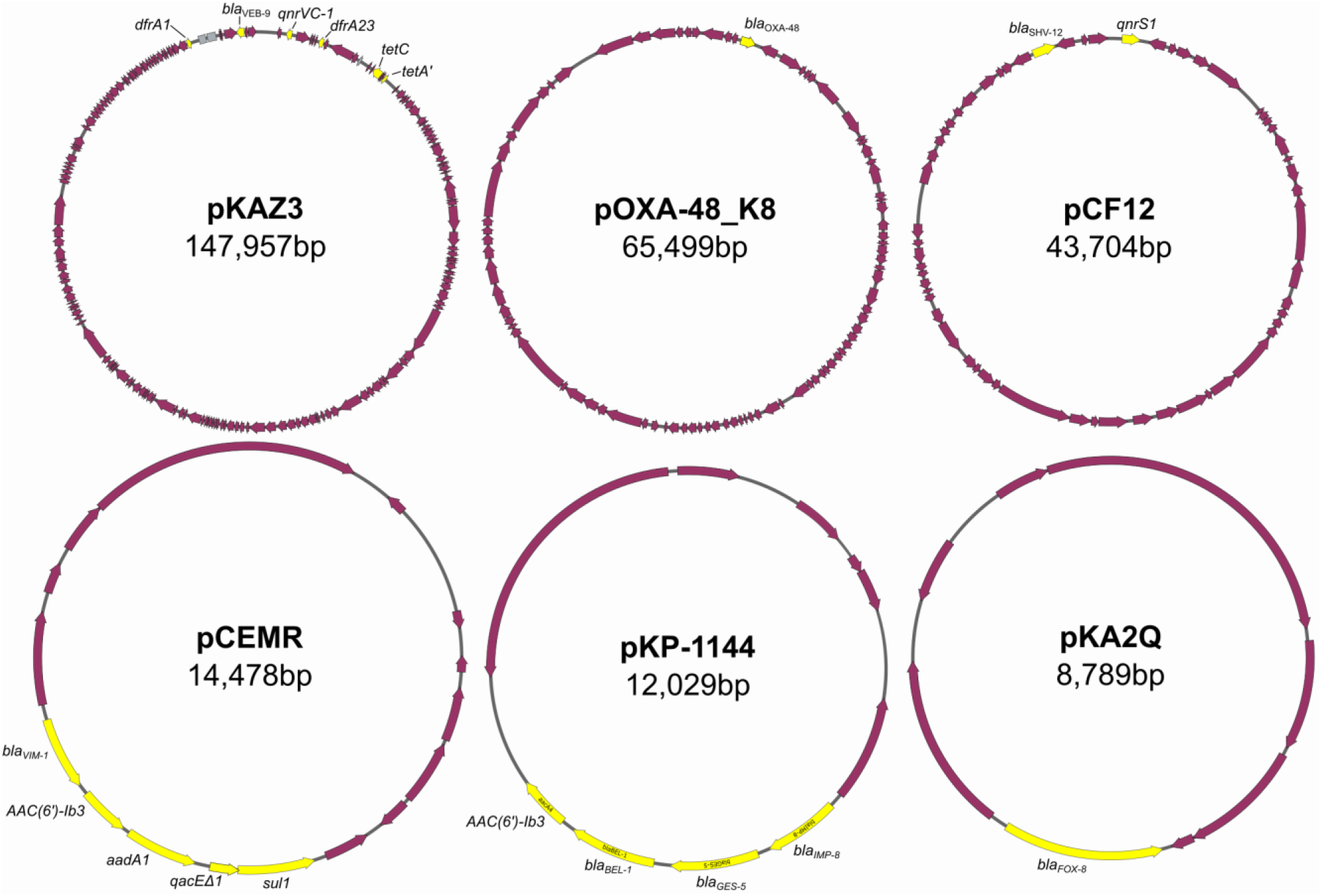
Plasmids used in this study. Scheme depicting the 6 plasmids used in this study. Antibiotic resistance genes are labeled and highlighted in yellow. Accession numbers are as follows: pKAZ3 (MT720905), pOXA-48_K8 (MT441554), pCF12 (MT720906), pCEMR (MT720903), pKP-1144 (MT720902), pKA2Q (MT720904). pOXA-48_K8 is a pOXA-48-like plasmid that shares a 100% coverage and >99% identity with the first described pOXA-48 plasmid (Poirel et al., 2012). For simplicity we refer to this plasmid as pOXA-48 throughout the text. Plasmids are not drawn to scale.

**Supplementary Figure 2.**
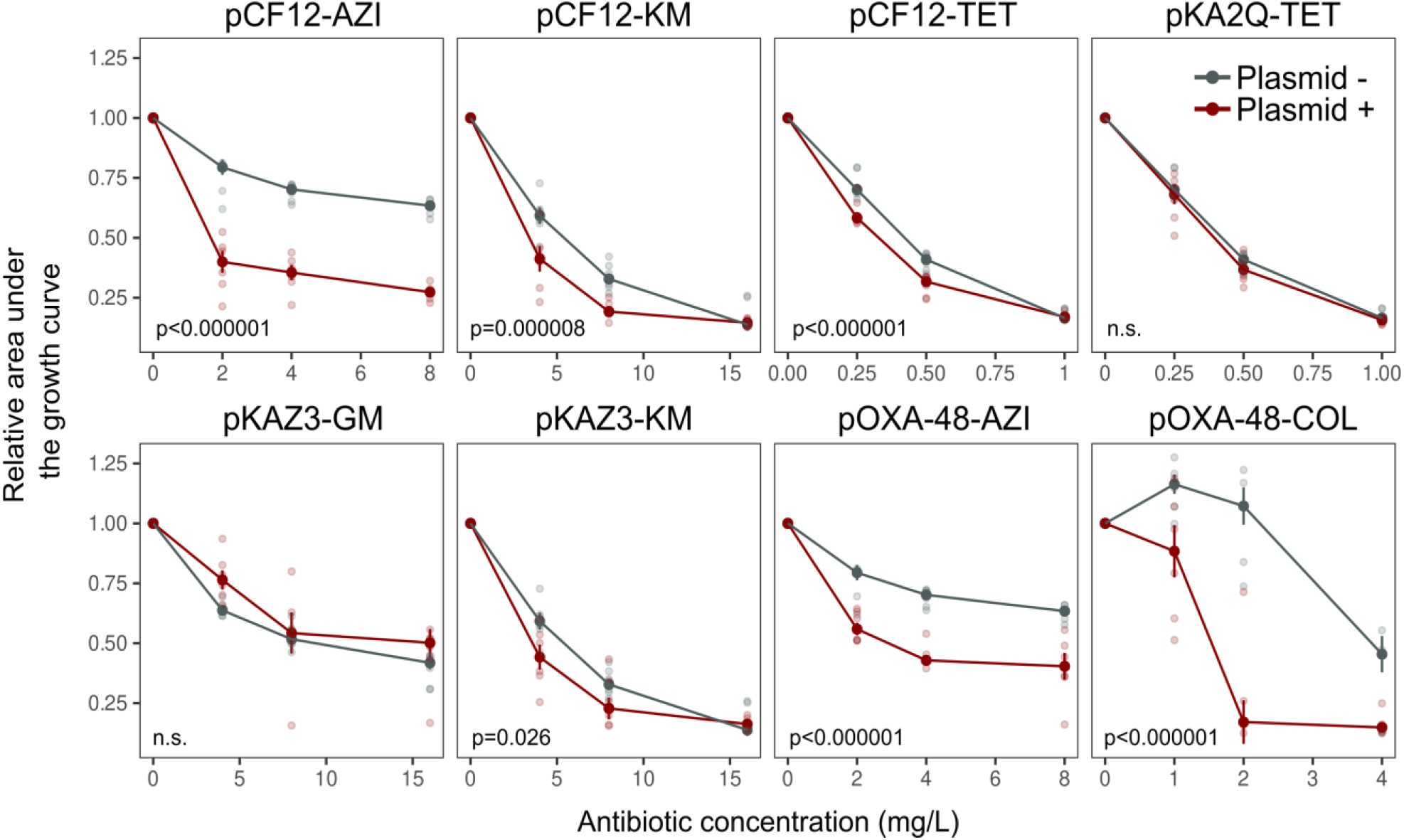
Analysis of bacterial growth curves. Comparison of the relative area under the growth curve of MG1655 (gray lines) and its plasmid-carrying derivatives (red) under different antibiotic concentrations. Each data point corresponds to the mean of 6-12 biological replicates and error bars correspond to the standard error of the mean. Individual data points are also represented. P-values are provided for the significantly different comparisons (ANOVA effect of ‘plasmid x antibiotic concentration’ interaction; ANOVA results are shown in Supplementary Table 2). AZI: azithromycin, COL: colistin, GM: gentamycin, KM: kanamycin, TET: tetracycline.

**Supplementary Table 1.** Antibiotic susceptibility results.

|  |  | Minimal inhibitory concentration (mg/L) |  |  |  |  |  |  |  |
| --- | --- | --- | --- | --- | --- | --- | --- | --- | --- |
| Drug class/target | Antibiotics | MG1655 |  | MG1655/<br>pOXA-48 | MG1655/<br>pKAZ3 | MG1655/<br>pKA2Q | MG1655/<br>pCF12 | MG1655/<br>pCEMR | MG1655/<br>pKP-M1144 |
|  |  | MH | LB |  |  |  |  |  |  |
| *B*-lactam/<br>Cell wall synthesis | Amoxicillin-clavulanic acid<br>(AMC) | 6 | 16 | >512 | 32 | 256 | 128 | 512 | 384 |
|  | Cefotaxime<br>(CTX) | 0.03 | 0.375 | 8 | 8 | 8 | 8 | 8 | 8 |
|  | Ceftazidime<br>(CAZ) | 0.5 | 0.375 | 0.5 | 32 | 32 | 32 | 32 | 32 |
|  | Ertapenem<br>(ERT) | 0.01 | 0.062 | 0.187 | 0.507 | 0.125 | 0.125 | 0.25 | 0.75 |
| Aminoglycoside/<br>Protein synthesis<br>(30S) | Gentamicin (GM) | 2 | 6 | 6 | 2 | 4 | 4 | 8 | 8 |
|  | Kanamycin<br>(KM) | 8 | 12 | 24 | 6 | 12 | 4 | 288 | 24 |
| Tetracycline/<br>Protein synthesis<br>(30S) | Tetracycline<br>(TET) | 2 | 4 | 4 | 32 | 2 | 2 | 4 | 4 |
|  | Tigecycline<br>(TGC) | 0.5 | 1 | 1 | 1 | 1 | 1 | 1 | 1 |
| Amphenicol/<br>Protein synthesis<br>(50S) | Chloramphenicol<br>(CM) | 8 | 8 | 16 | 8 | 8 | 8 | 8 | 8 |
| Macrolide/<br>Protein synthesis<br>(50S) | Azithromycin<br>(AZI) | 16 | 16 | 8 | 12 | 12 | 8 | 12 | 12 |
| Quinolone/ | Ciprofloxacin | 0.5 | 0.5 | 0.75 | 8 | 1 | 8 | 2 | 3 |
| DNA gyrase | (CIP) |  |  |  |  |  |  |  |  |
| Rifamycin/<br>RNA polymerase | Rifampicin (RIF) | 8 | 8 | 8 | 24 | 24 | 8 | 16 | 256 |
| Polymyxin/<br>Lipopolysaccharide | Colistin (COL) | 4 | 8 | 4 | 6 | 8 | 6 | 8 | 6 |

MIC of the 13 antibiotics for *E. coli* MG1655 in LB and MH medium and the MIC of the six plasmid-carrying MG1655 strains in LB medium. Median value of 4-5 independent biological replicates.

**Supplementary Table 2.**
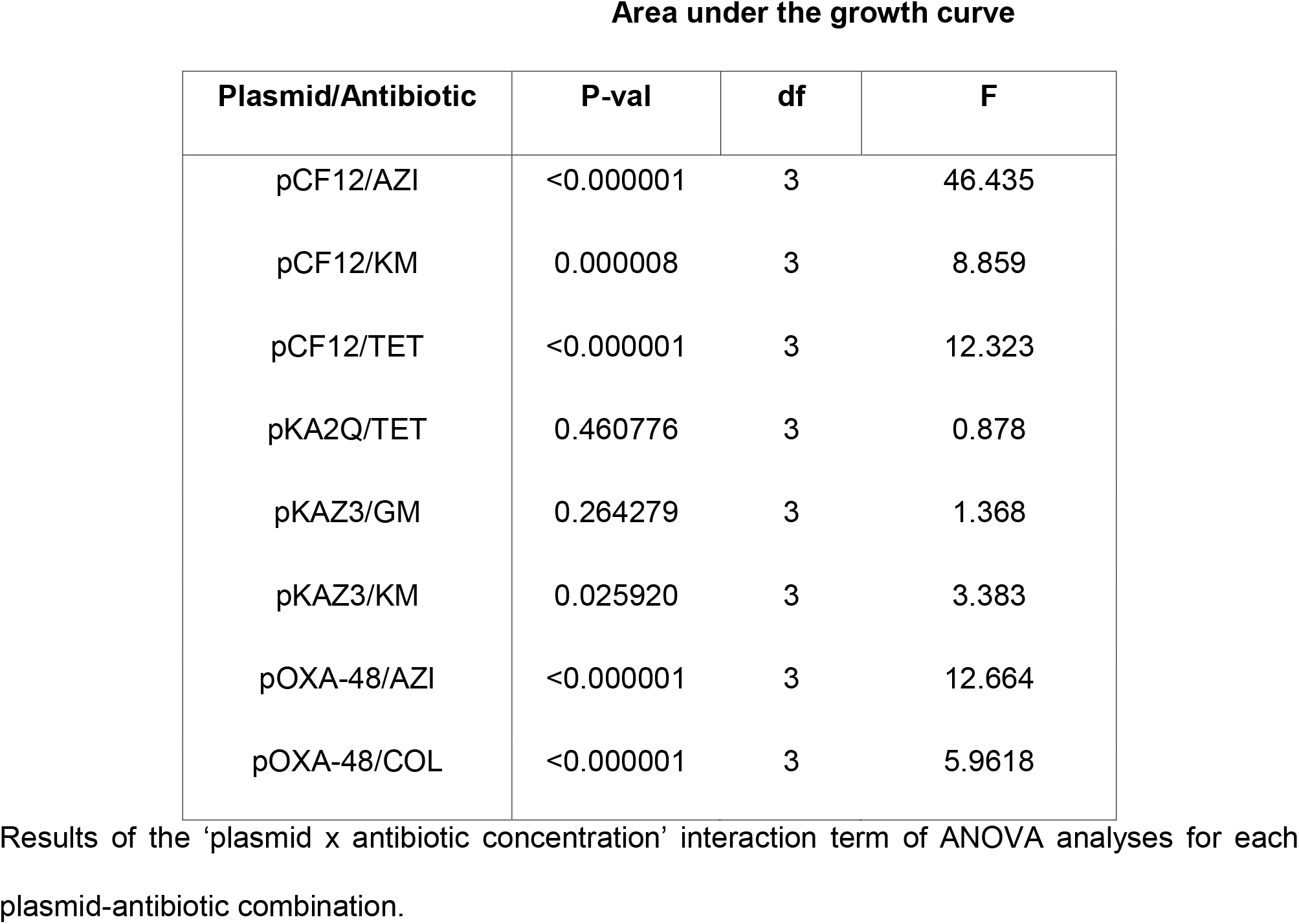
ANOVA results of the growth curves of plasmids producing CS effects.

**Supplementary Table 3.** Chromosomal mutations accumulated during plasmid acquisition.

| Strain | Gene | Gene product | Type | Position<sup>A</sup> | Reference<sup>B</sup> | Alternative | Effect |
| --- | --- | --- | --- | --- | --- | --- | --- |
| MG1655/pKP-M1144 | *gadB* | Glutamate decarboxylase beta | SNP | 642/1401 | G | A | Synonymous (Thr214Thr) |
| MG1655/pKP-M1144 | *selB* | Selenocysteine-specific elongation factor | SNP | 1256/1845 | G | T | Missense (Trp419Leu) |
| MG1655/pCEMR | *ppk* | Polyphosphate kinase | SNP | 1488/2067 | T | C | Synonymous (Arg496Arg) |
| MG1655/pCEMR | *selB* | Selenocysteine-specific elongation factor | SNP | 1256/1845 | G | T | Missense (Trp419Leu) |
| MG1655/pKA2Q | *cpxA* | Sensor histidine kinase | SNP | 515/1374 | A | G | Missense (Met172Thr) |
| MG1655/pCF12 | Intergenic | - | Insertion | 1,212,080 | AGGGGGGGGGT | AGGGGGGGGGGT | - |

<sup>A</sup>Position relative to the start of the open reading frame, except for the intergenic mutation which refers to the chromosomal coordinate. <sup>B</sup>Reference genome is plasmid-free MG1655 (deposited in Sequence Read Archive [SRA], BioProject ID: PRJNA644278)

**Supplementary Table 4.** Characteristics of the *E. coli* clinical isolates used in this study.

| Strain | Phylogroup | Sequence Type | Plasmids | Resistance genes |
| --- | --- | --- | --- | --- |
| Ec02 | B1 | 453 | Col-like, IncFIB, IncI, IncFIC, IncY | *aadA5*, *blaCTX-M-1*, *blaTEM-1B*, *dfrA17*, *sul2*, *tet(B)*, *mdf(A)* |
| Ec03 | B2 | 131 | Col-like, IncFIB, IncFII, IncB/O/K/Z | *blaCTX-M-14*, *blaTEM-1B*, *mdf(A)* |
| Ec04 | B2 | 131 | Col-like, IncFIB, IncFII, IncFIA | *aadA5*, *aph(3'')-Ib*, *aph(6)-Id*, *blaCTX-M-14*, *blaTEM-1B*, *dfrA17*, *mph(A)*, *sul1*, *sul2*, *tet(A)*, *mdf(A)* |
| Ec06 | A | 10 | Col-like, IncFII, IncI | *aadA5*, *blaCTX-M-1*, *sul2*, *tet(A)*, *mdf(A)* |
| Ec10 | D | 69 | Col-like, IncFII, pEC4115 | *aac(3)-Iva*, *aph(3'')-Ib*, *aph(4)-Ia*, *aph(6)-Id*, *blaCTX-M-1*, *mph(A)*, *mdf(A)* |
| Ec18 | F | 648 | Col-like, IncY | *aadA5*, *blaCTX-M-14*, *dfrA17*, *mph(A)*, *sul1*, *mdf(A)* |
| Ec20 | B2 | 131 | IncFIB, IncI, IncFIA, IncFIC | *aac(3)-Iid*, *blaCTX-M-15*, *blaTEM-1B*, *mdf(A)* |
| Ec21 | D | 405 | Col-like, IncFIB, IncFII, IncFIA, p0111 | *aac(3)-Iia*, *aph(3'')-Ib*, *aph(6)-Id*, *blaCTX-M-15*, *floR*, *sul2*, *tet(A)*, *tet(B)*, *mdf(A)* |
| Ec25 | A | 10 | IncFIB, IncFII, IncX, IncFIC | *aph(3')-Ia*, *aph(6)-Id*, *blaCTX-M-3*, *dfrA14*, *fosA3*, *sul2*, *mdf(A)* |

## Notes

### Competing Interest Statement

The authors have declared no competing interest.

